# Genome wide association study of hippocampal subfield volume in PTSD cases and trauma-exposed controls

**DOI:** 10.1101/456988

**Authors:** Rajendra A. Morey, Melanie E. Garrett, Jennifer S. Stevens, Emily Clarke, Courtney C. Haswell, Sanne J.H. van Rooij, Negar Fani, Adriana Lori, VA Mid-Atlantic MIRECC Workgroup, Christine E. Marx, Jean C. Beckham, Gregory McCarthy, Michael A. Hauser, Allison E. Ashley-Koch

## Abstract

Behavioral, structural, and functional neuroimaging have implicated the hippocampus as a critical brain region in PTSD pathogenesis. We conducted a GWAS of hippocampal subfield volumes in a sample of recent military veteran trauma survivors (n=157), including some with PTSD (n=66). Covariates in our analysis included lifetime PTSD diagnosis, sex, intracranial volume, genomic estimates of ancestry, and childhood trauma. Interactions between genetic variants and lifetime PTSD or childhood trauma were interrogated for SNPs with significant main effects. Several genetic associations surpassed correction for multiple testing for several hippocampal subfields, including fimbria, subiculum, cornu ammonis-1(CA1), and hippocampal amygdala transition area (HATA). One association replicated in an independent cohort of civilians with PTSD (rs12880795 in *TUNAR* with L-HATA volume, *p*=3.43 × 10^-7^ in the discovery and *p*=0.0004 in the replication cohort). However, the most significant association in the discovery data set was between rs6906714 in *LINC02571* and R-fimbria volume (*p*=5.99 ×10^-8^, *q*=0.0056). Interestingly, the effect of rs6906714 on R-fimbria volume increased with childhood trauma (G*E interaction *p*=0.022). In addition to variants in long intergenic non-coding RNAs (lincRNAs), we identified SNPs associated with hippocampal subfield volume, which are also quantitative trait loci (QTLs) for genes involved in RNA editing of glutamate receptor subunits (GluRs), oxidative stress, and autoimmune disorders. Genomic regions, some with putative regulatory roles, influence the volume of hippocampal subfields. Neuroimaging phenotypes may offer important insight into the genetic architecture and neurobiological pathways relevant to PTSD, as well as in the identification of potential biomarkers and drug targets for PTSD.

## INTRODUCTION

Posttraumatic stress disorder (PTSD), which affects about 8% of the US population, can develop following exposure to trauma and manifests with the hallmark symptoms of hyperarousal, re-experiencing, and avoidance. Importantly, deficits in memory, including declarative memory, fragmented autobiographical or trauma-related memories and amnesia about the details of trauma have been demonstrated in PTSD(1). The hippocampus has long been implicated in PTSD because of its role in memory formation and retrieval(2), as well as the observation of lower hippocampal volume in individuals with PTSD(3, 4). It is unclear whether lower hippocampal volume is a consequence of developing PTSD or occurs as the result of heretofore unknown genetic or biological vulnerabilities to developing PTSD. The genetic modulators of hippocampal volume have been established in non-clinical populations with genome wide association studies(GWAS)(5-7). While the SNP-based heritability for hippocampal subfields is modest in normative samples(*h^2^* = 0.14 to 0.27), the specific genetic loci associated with hippocampal subfield volume in PTSD have not been investigated.

There are several advantages to the exploration of neuroimaging phenotypes over diagnostic phenotypes, which consist of a constellation of symptoms or behaviors. The first advantage is the enhanced accuracy of imaging phenotypes over a subjectively assessed diagnostic phenotype. This may eliminate a significant component of noise from the overall model. Second, the imaging phenotype is likely to be closer to the action of genes than the diagnostic phenotype, which is determined by a compilation of several structural and functional components. A frequent rationale for imaging genetics studies compared to diagnostic phenotypes has been larger effect sizes, but this not been borne out by data thus far. Nonetheless, twin-based heritability estimates of 30-35% obtained from GWAS of PTSD(8) are dramatically smaller than the twin-based heritability estimates of hippocampal subfields(56% for hippocampal fissure; 67% for the hippocampal amygdala transition area(HATA); 84% for CA1)(9). The SNP heritability estimates of PTSD in females is 29%, but could not be distinguished from zero in males(10).

Subfields of the hippocampus are involved in discrete aspects of memory encoding and consolidation. For example, the dentate gyrus(DG) is important in *pattern separation*, a process where salient features of memories are contrasted in order to distinguish similar but discrete events(11, 12). By contrast, the entorhinal cortex(EC) and cornu ammonis subfield-3(CA3) are crucial to *pattern completion*, which involves recognizing overlapping features of events. Pattern completion is a widely-investigated model of PTSD given its important implication in contextual fear conditioning(13, 14). Preclinical animal models show that CA1 is involved in context-specific memory retrieval following extinction(15). As such, the CA1 hippocampal subfield has been implicated strongly in conditioned fear and its extinction(16). The deficits in episodic memory, contextual memory, and extinction failure suggest that CA1, CA3, and dentate subfields may play a role in developing PTSD(17).

Two GWAS from the ENIGMA Consortium have examined effects on overall hippocampal volume with significant results at the genome-wide level. Stein et al(7) identified and replicated two quantitative trait loci for hippocampal volumes in a large sample that included non-clinical and clinical samples with neuropsychiatric diagnoses. The NORMENT Center recently reported on 21,297 samples used to identify 15 unique genomewide significant loci across six hippocampal subfields, of which eight loci had not been previously linked to the hippocampus(5). The SNPs mapped to genes associated with neuronal differentiation, locomotor behavior, schizophrenia, and Alzheimer disease. These studies have demonstrated the ability to detect genetic associations with neuroimaging phenotypes that may be relevant to PTSD. However, as yet, there have been no published studies that have examined genetic predictors of hippocampal subfields in PTSD. The present study examines two samples of highly traumatized individuals. The discovery sample of US military veterans with roughly equal number of European- and African-Americans experienced high levels of trauma in the military and in some cases during childhood and adolescence(18). The replication sample of civilians consists of African-American women in Atlanta [Georgia USA] who experienced high rates of sustained trauma and interpersonal violence. Based on the role of the dentate gyrus in pattern separation, CA3 in pattern completion, and CA1 in fear extinction, we hypothesized the presence of genetic variants that influence subfield volumes, which are obtained from segmentation of high-resolution structural MRI, in these enriched samples of trauma exposed individuals with and without PTSD.

## METHODS

### Participants and Clinical Measures

#### Discovery Cohort

The discovery cohort consisted of 157 participants from a repository [Mid-Atlantic MIRECC, Durham NC] of Iraq and Afghanistan era military service members who contributed blood for genotyping, clinical assessment data, and MRI scans. Participants were screened for inclusion/exclusion criteria based on information available in the repository. Important exclusion criteria included presence of psychotic symptoms, high risk of suicide, contraindication to MRI, current substance abuse, neurological disorders, and age over 65 years. To reduce the effects of population stratification in a multi-racial sample, analyses were limited to non-Hispanic black(NHB;n=74) and non-Hispanic white(NHW;n=83) participants from these studies who consented to the genetic and imaging components and had data available at the time of analysis. All participants provided written informed consent to participate in procedures reviewed and approved by the Institutional Review Boards at Duke University and the Durham VA Medical Center. Participants were evaluated for PTSD, trauma exposure, and other psychiatric comorbidities as described in the Supplementary Materials.

#### Replication Cohort

The replication sample consisted of 133 participants drawn from the Grady Trauma Project(GTP), which is a large study of PTSD risk factors that recruited participants from a low-socioeconomic status urban cohort of outpatients in general medical clinics at Grady Hospital [Atlanta, GA]. To minimize heterogeneity, the sample was restricted to African-American women. Seventy-nine participants met criteria for diagnosis of PTSD, and 54 trauma-exposed controls(TC) did not. Study procedures were approved by the Institutional Review Board of Emory University and the Research Oversight Committee of Grady Memorial Hospital, and all participants provided written informed consent prior to participating. Participants were evaluated for PTSD, trauma exposure, and other psychiatric comorbidities as described in the Supplementary Materials.

### MRI acquisition in discovery sample

#### Discovery sample

Images were acquired on a General Electric 3-Tesla Signa EXCITE scanner equipped with an 8-channel head coil. High-resolution T1-weighted whole-brain images using 3D-FSPGR were acquired axially(TR/TE/flip angle=7.484 ms/2.984 ms/12°, FOV=256 mm, 1-mm slice thickness, 166 slices, 1-mm^3^ voxel size, 1 excitation).

#### Replication sample

Scanning took place on a 3.0 T Siemens Trio with echo-planar imaging. High-resolution T1-weighted whole-brain anatomical scans were collected using a 3D MP-RAGE sequence, with 176 contiguous 1-mm sagittal slices(TR/TE/TI = 2000/3.02/900 ms, 1-mm^3^ voxel size).

### Hippocampal Subfield Volume Analysis

Identical procedures were applied to the discovery and replication sample to perform automated hippocampal subfield segmentation using FreeSurfer 6.0.0, which were previously reported in detail(17) and partially recapitulated in the Supplementary Materials.

### Quality Control

Identical visual inspection procedures were applied to all T1 images from the discovery(RAM, CCH) and replication cohorts(JSS, SVR) for quality assurance purposes. We applied quality assurance for hippocampal subfield segmentations using ENIGMA-PTSD protocols reported in(17). All participants passed both rounds of quality control.

### Genotyping

#### Discovery Sample

GWAS data were generated as part of a larger parent study of 2,312 PTSD cases and controls, as previously described(19) and recapitulated in the Supplementary Materials.

#### Replication sample

Saliva collected in Oragene vials underwent DNA extraction followed by genotyping on Illumina platforms and QC measures with PLINK. Levels of heterozygosity, devioation from Hardy-Weinberg, screening for relatedness, and low call rates were part of QC(Supplementary Materials).

### Statistical Analysis

Principal components analysis(PCA) was run using the *smartpca* program from the software package EIGENSOFT(20) to assess population stratification in the discovery sample. One principal component was necessary to account for the population variability observed in this subset of individuals, essentially distinguishing the NHB from the NHW subjects. Linear regression was utilized using PLINK(21) to test for association between subcortical volume and each SNP, assuming an additive genetic model. Left and right brain hemisphere volumes were analyzed separately. All subcortical volumes were normally distributed except the left fimbria, which was log transformed for analysis. Covariates included sex, age, one population stratification principal component, lifetime PTSD diagnosis, intracranial volume, and childhood trauma(0=no childhood trauma; 1=exposure to a single category of childhood trauma; 2=exposure to two or more categories of childhood trauma, as reported from TLEQ items 12,13,15,16, and 17). To reduce genetic redundancy in this imputed dataset, we used the linkage disequilibrium(LD) clumping method in PLINK, choosing an r^2^ threshold of 0.25 and a 500kb window, as reported previously(22). False discovery rate(FDR) q-values were generated using PROC MULTTEST in SAS version 9.4. The FDR correction was applied only across SNPs but not across the 12 hippocampal subfields and two hemispheres because the inherent correlation between subfield volumes would lead to overly conservative inferences. The lack of correction for the number of subfields yields somewhat permissive inferences in this initial exploration which motivated the identification of an independent replication data set. Post-hoc interaction analyses were performed only for the SNPs with significant main effects. SNP*childhood trauma and SNP*lifetime PTSD interactions were investigated for associations with subfield volumes.. Manhattan plots and Q-Q plots were produced using the R package *qqman*(23) and interaction plots were generated using the R package *ggplot2*(24).

In the replication dataset, PLINK was used to prune the autosomal data in windows of 50 bp, removing 1 SNP from each pair of SNPs with r^2^>0.05 to obtain a set of roughly independent markers(~50 000 SNPs) for use in PCA. Subjects who fell within 3 standard deviations of the medians of the first and second PCs were retained. Linear regression was performed in the replication sample using the same software and model described above for the discovery cohort.

## RESULTS

Important clinical and sociodemographic information by cohort is reported in **Table 1**. Compared to the discovery cohort, the replication cohort comprises a higher percentage of females(p<0.0001), African-Americans(p<0.0001), subjects reporting alcohol misuse(p=0.0379), and lifetime PTSD(p=0.0032). Average age and distribution of childhood trauma did not significantly differ between the two cohorts.

**Table 1.**
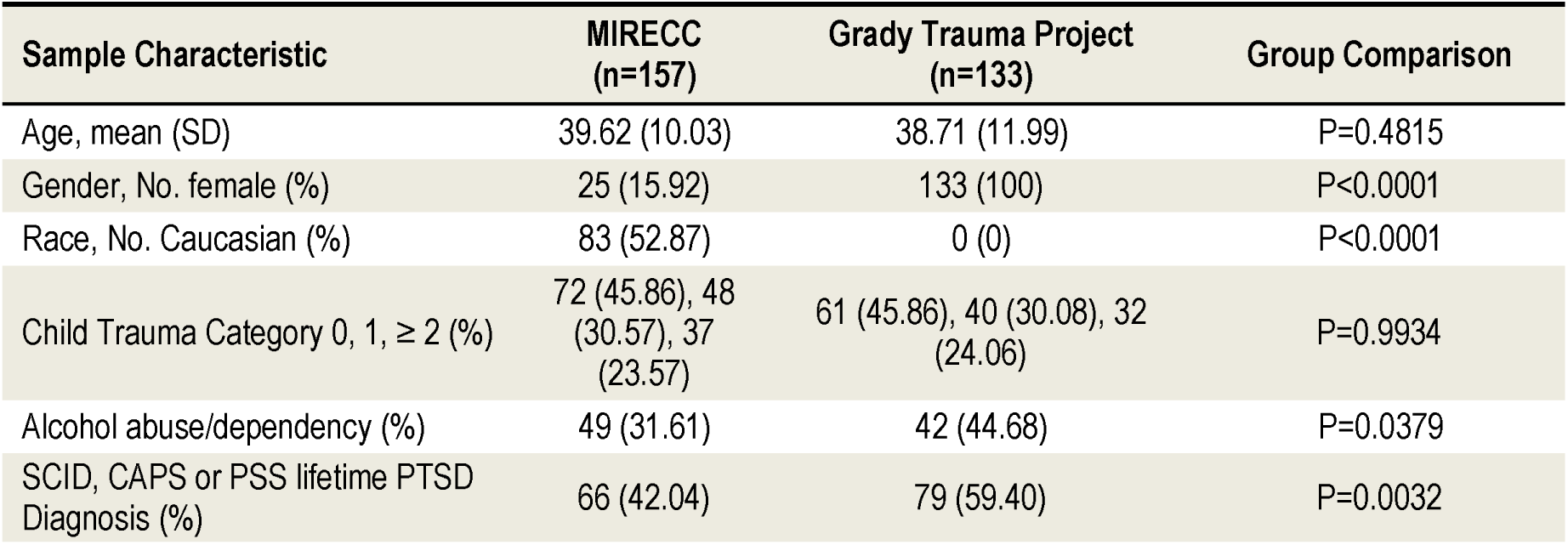
Demographic and Clinical Information § The Davidson Trauma Scale was used in lieu of the CAPS for 5 subjects for which CAPS was unavailabel. Abbreviations: SD=standard deviation, No=number, AUDIT=Alcohol Use Disorders Identification Test, SCID-IV=Structured Clinical Interview for DSM-IV, CAPS-IV Clinician Administered PTSD Scaler with DSM-IV criteria.

| Sample Characteristic | MIRECC<br>(n=157) | Grady Trauma Project<br>(n=133) | Group Comparison |
| --- | --- | --- | --- |
| Age, mean (SD) | 39.62 (10.03) | 38.71 (11.99) | P=0.4815 |
| Gender, No. female (%) | 25 (15.92) | 133 (100) | P<0.0001 |
| Race, No. Caucasian (%) | 83 (52.87) | 0 (0) | P<0.0001 |
| Child Trauma Category 0, 1, ≥ 2 (%) | 72 (45.86), 48<br>(30.57), 37<br>(23.57) | 61 (45.86), 40 (30.08), 32<br>(24.06) | P=0.9934 |
| Alcohol abuse/dependency (%) | 49 (31.61) | 42 (44.68) | P=0.0379 |
| SCID, CAPS or PSS lifetime PTSD<br>Diagnosis (%) | 66 (42.04) | 79 (59.40) | P=0.0032 |

We identified several SNPs associated with different hippocampal subfield volumes that survived correction for multiple testing(**Table 2**) in the discovery cohort. The most significant associations(q<0.01) involved R-fimbria volume(**Figure 1**). The SNP rs6906714, located in *LINC02571* on chromosome 6(**Supplementary Figure 1**), was associated with R-fimbria volume(p=5.99 × 10^-8^, q=0.0056), such that for each additional G allele, R-fimbria volume increased by 15.73 mm^3^. An intergenic SNP, rs17012755 on chromosome 2, was also associated with R-fimbria volume(p=6.05 x 10 ^-8^, q=0.0056), Each additional copy of the A allele increased R-fimbria volume by 22.01 mm^3^. In addition to the main effects, we also identified an interaction between rs6906714 and childhood trauma affecting R-fimbria volume(p=0.022, **Figure 2**). As exposure to childhood trauma increased, the effect of rs6906714 genotype on R-fimbria volume became stronger. Specifically, among individuals who did not experience any childhood trauma, the association between rs6909714 genotype and R-fimbria was modest(p=0.037; beta=9.272 mm^3^). Among those who did experience childhood trauma(either 1 or 2+ categories), the association was more robust with *p*=0.0002; beta=20.19 mm^3^ for 1 category of childhood trauma and *p*=0.0002; beta=25.19 mm^3^ for 2+ categories. There was no appreciable difference in the effect of rs6906714 genotype on R-fimbria volume among those who experienced 2+ categories of childhood trauma compared to those who experienced a single category of childhood trauma. Several additional significant main effects were observed(q<0.05) in the discovery cohort with different hippocampal subfields, some intergenic and some residing in genes. The SNP rs7196581 in *RBBP6* was associated with two different subfields: L-CA1(p=1.51 x 10^-7^, q=0.0280) and R-HATA(p=4.58 x 10^-7^, q=0.0424), such that for each additional G allele, L-CA1 volume increased by 53.62 mm^3^ and R-HATA volume increased by 7.94 mm^3^. Also associated with increased R-HATA volume was the T allele of rs73008928 in *ARMC6*(p=2.33 × 10^-7^, q=0.0424). Three significant associations were identified with L-HATA volume: rs12880795 in *TUNAR*(p=3.43 × 10^-7^, q=0.0372), rs2553974 in *LOC105376580*(p=4.02 × 10^-7^, q=0.0372), and rs3811492 in *LINC01736*(p=8.13 x 10^-7^, q=0.0498). Finally, two intergenic SNPs were associated with L-fimbria and R-subiculum volumes: rs76832471 on chromosome 2(p=6.51× 10^-8^, q=0.0121) and rs9499406 on chromosome 6(p=8.19 × 10^-8^, q=0.0151), respectively.

**Figure 1a.**
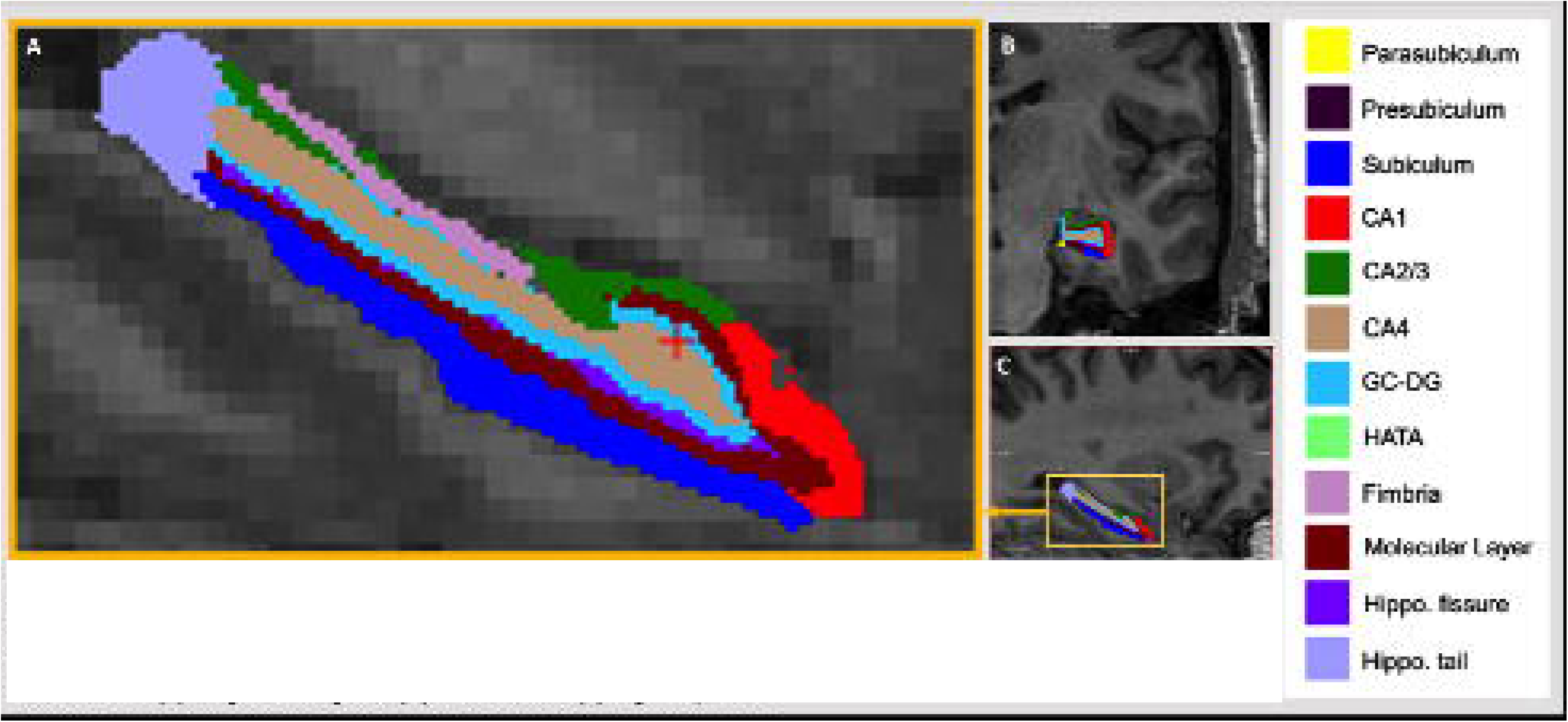
Automated segmentation of the hippocampus into 12 subfields in each hemisphere of the brain was performed with FreeSurfer vE.0 [beta version]. Subfield Images of CA1, CA2/3, CA4, dentate gyrus, hippocampal-amygdala transition area |HATA|, subiculum, tail, fissure, presubiculum, parasubiculum, molecular layer, fimbria are shown in **[a]** magnified signal. **[b]** cononal, and **[c]** sagictal planes.

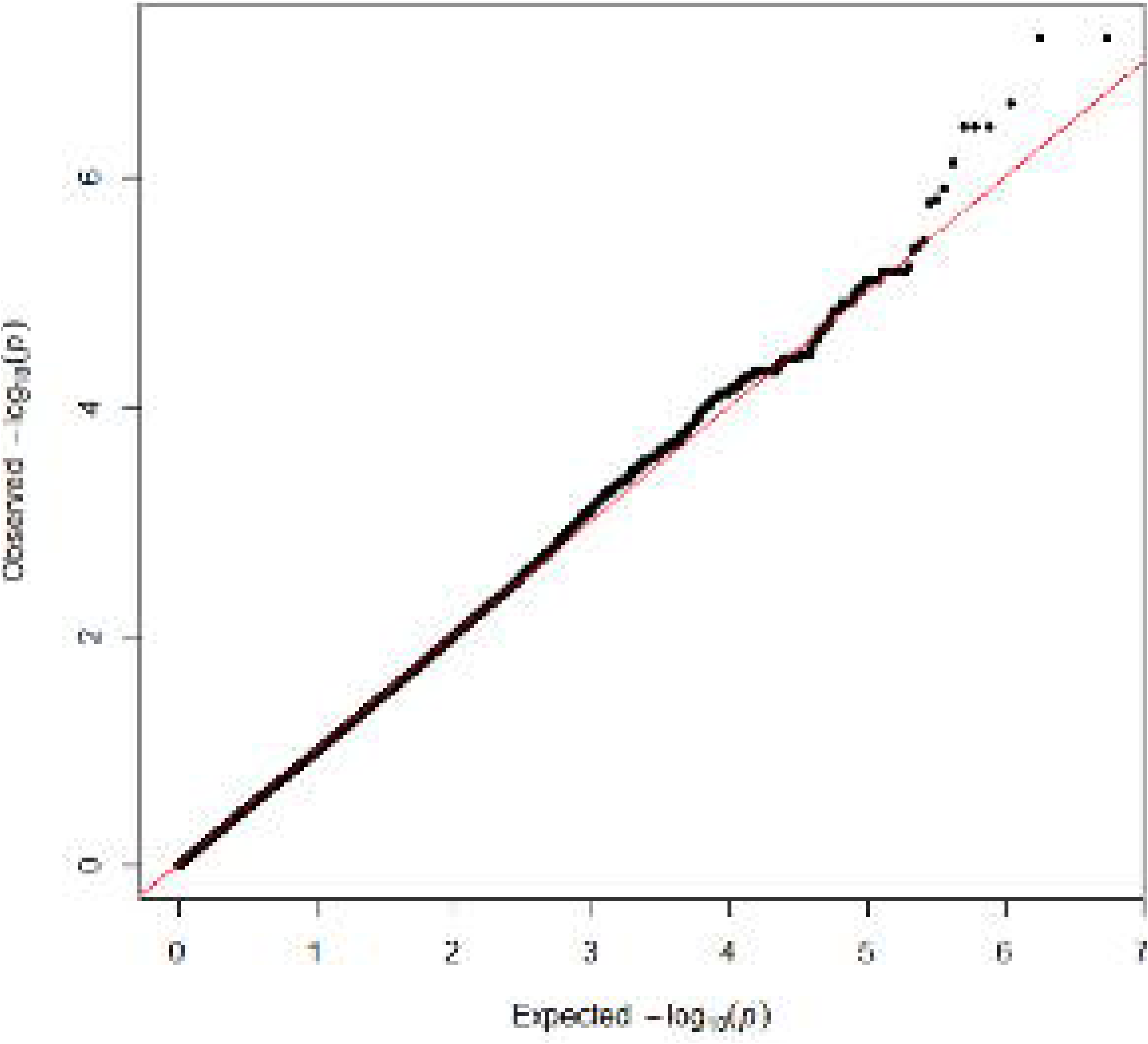

**Table 2.**
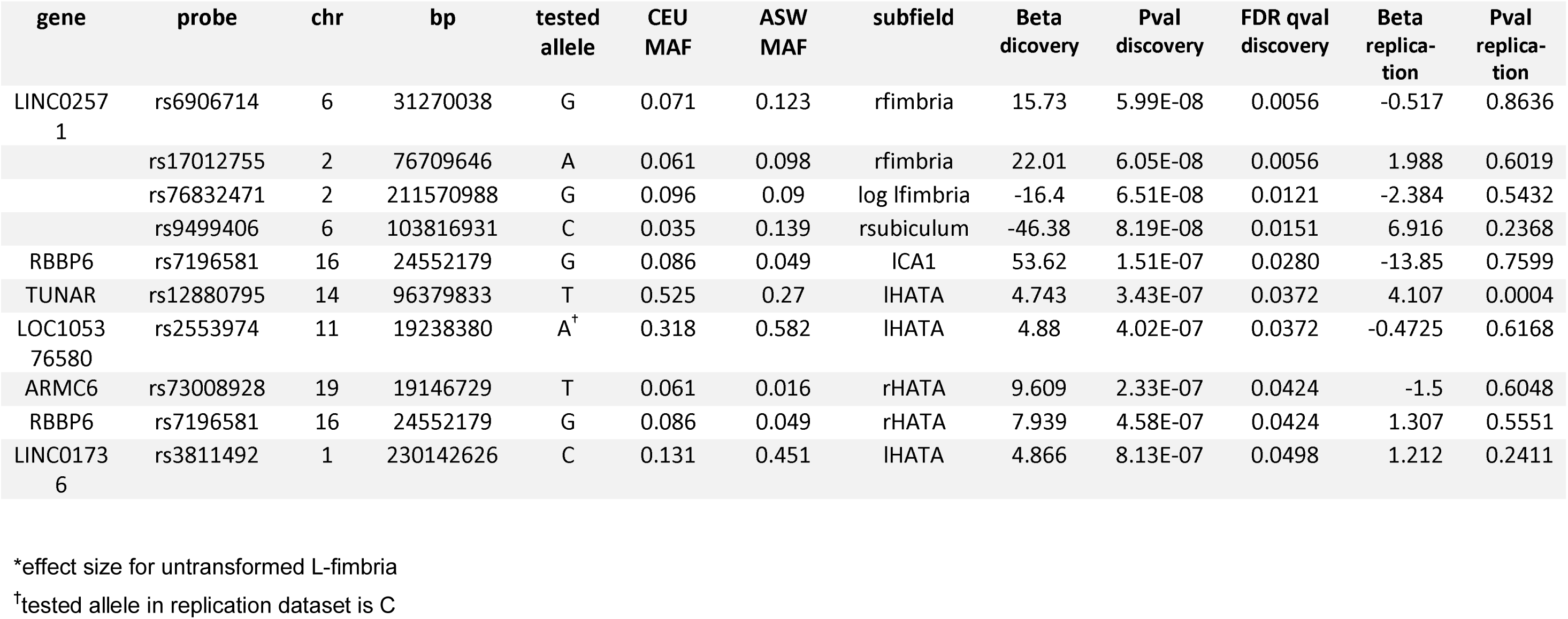
FDR significant main effects of genetic variants on hippocampal subfield volume.

| gene | probe | chr | bp | tested allele | CEU MAF | ASW MAF | subfield | Beta discovery | Pval discovery | FDR qval discovery | Beta replication | Pval replication |
| --- | --- | --- | --- | --- | --- | --- | --- | --- | --- | --- | --- | --- |
| LINC0257<br>1 | rs6906714 | 6 | 31270038 | G | 0.071 | 0.123 | rfimbria | 15.73 | 5.99E-08 | 0.0056 | -0.517 | 0.8636 |
|  | rs17012755 | 2 | 76709646 | A | 0.061 | 0.098 | rfimbria | 22.01 | 6.05E-08 | 0.0056 | 1.988 | 0.6019 |
|  | rs76832471 | 2 | 211570988 | G | 0.096 | 0.09 | log lfimbria | -16.4 | 6.51E-08 | 0.0121 | -2.384 | 0.5432 |
|  | rs9499406 | 6 | 103816931 | C | 0.035 | 0.139 | rsubiculum | -46.38 | 8.19E-08 | 0.0151 | 6.916 | 0.2368 |
| RBBP6 | rs7196581 | 16 | 24552179 | G | 0.086 | 0.049 | ICA1 | 53.62 | 1.51E-07 | 0.0280 | -13.85 | 0.7599 |
| TUNAR | rs12880795 | 14 | 96379833 | T | 0.525 | 0.27 | IHATA | 4.743 | 3.43E-07 | 0.0372 | 4.107 | 0.0004 |
| LOC1053<br>76580 | rs2553974 | 11 | 19238380 | A<sup>†</sup> | 0.318 | 0.582 | IHATA | 4.88 | 4.02E-07 | 0.0372 | -0.4725 | 0.6168 |
| ARMC6 | rs73008928 | 19 | 19146729 | T | 0.061 | 0.016 | rHATA | 9.609 | 2.33E-07 | 0.0424 | -1.5 | 0.6048 |
| RBBP6 | rs7196581 | 16 | 24552179 | G | 0.086 | 0.049 | rHATA | 7.939 | 4.58E-07 | 0.0424 | 1.307 | 0.5551 |
| LINC0173<br>6 | rs3811492 | 1 | 230142626 | C | 0.131 | 0.451 | IHATA | 4.866 | 8.13E-07 | 0.0498 | 1.212 | 0.2411 |

\*effect size for untransformed L-fimbria
<sup>†</sup>tested allele in replication dataset is C

Aside from the significant interaction described above between rs6906714 and childhood trauma on R-fimbria volume, no other SNP displaying a significant main effect demonstrated a significant interaction with either childhood trauma or lifetime PTSD diagnosis and hippocampal subfield volume in the discovery cohort.

The ten significant main effects identified in the discovery cohort(q<0.05) were tested for significance in the replication cohort. Despite significant differences in the composition of discovery and replication cohorts(**Table 1**), we observed an association between rs12880795 genotype and L-HATA volume(p=0.0004) in the replication cohort. No other main effect of SNP on hippocampal subfield volume observed in the discovery cohort was replicated. However, we did detect a significant interaction between rs6906714 and lifetime PTSD on R-fimbria volume in the replication cohort(p=0.0421; **Figure 2**). Similarly, while we found no main effect of rs17012755 on R-fimbria volume in the discovery cohort, we observed a significant interaction between rs17012755 and lifetime PTSD on R-fimbria volume in the replication cohort(p=0.0219; **Figure 3**).

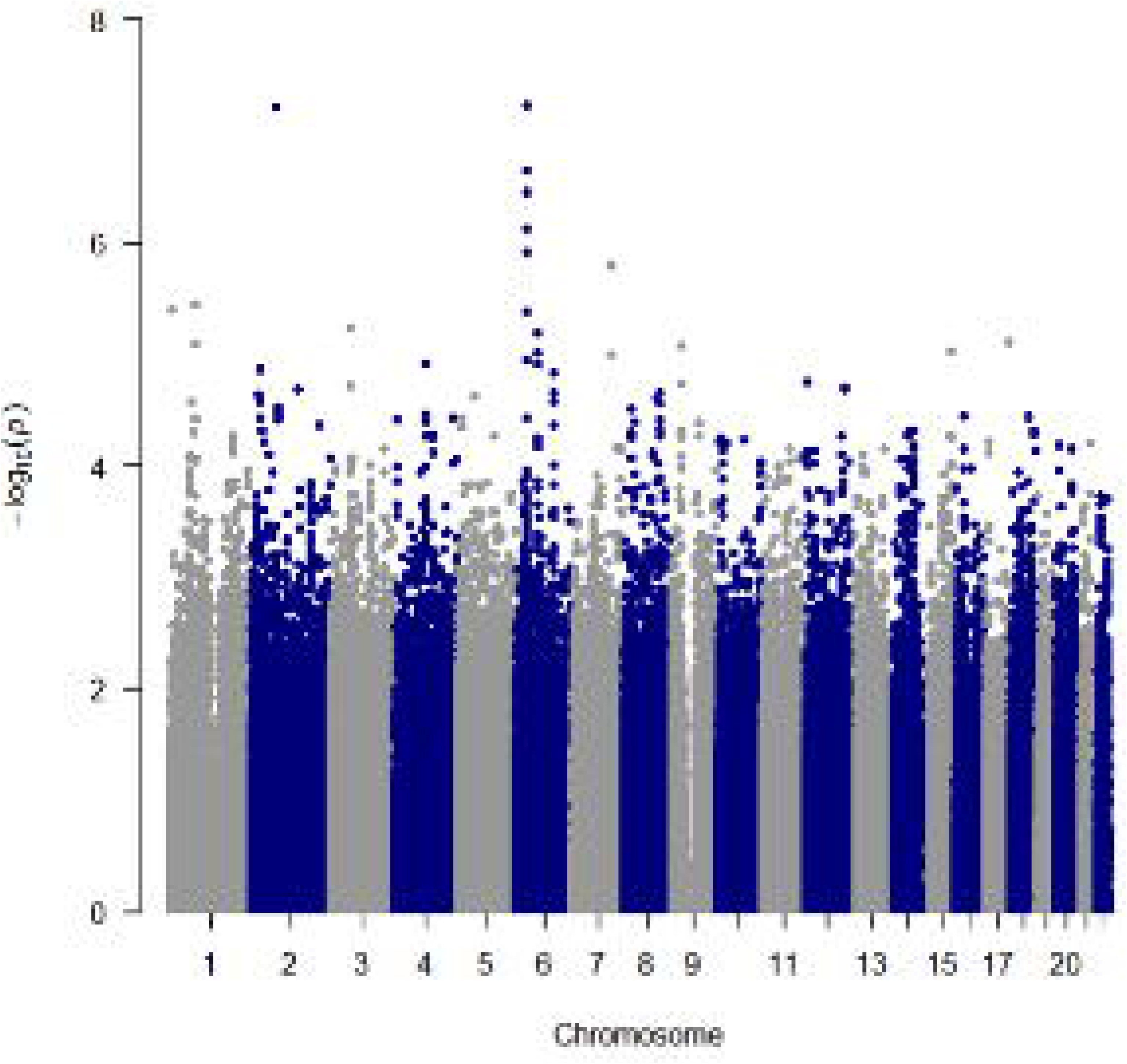

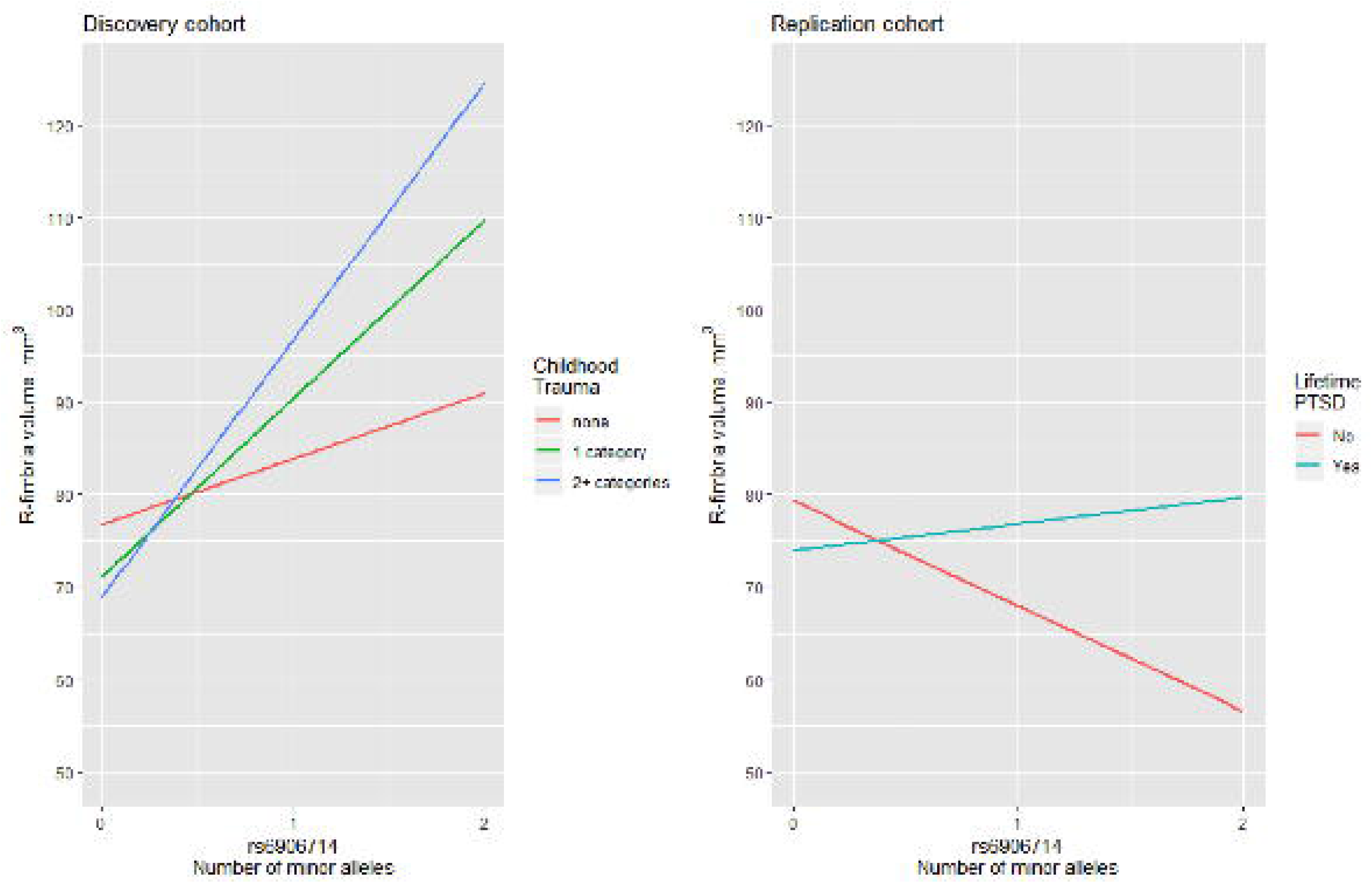

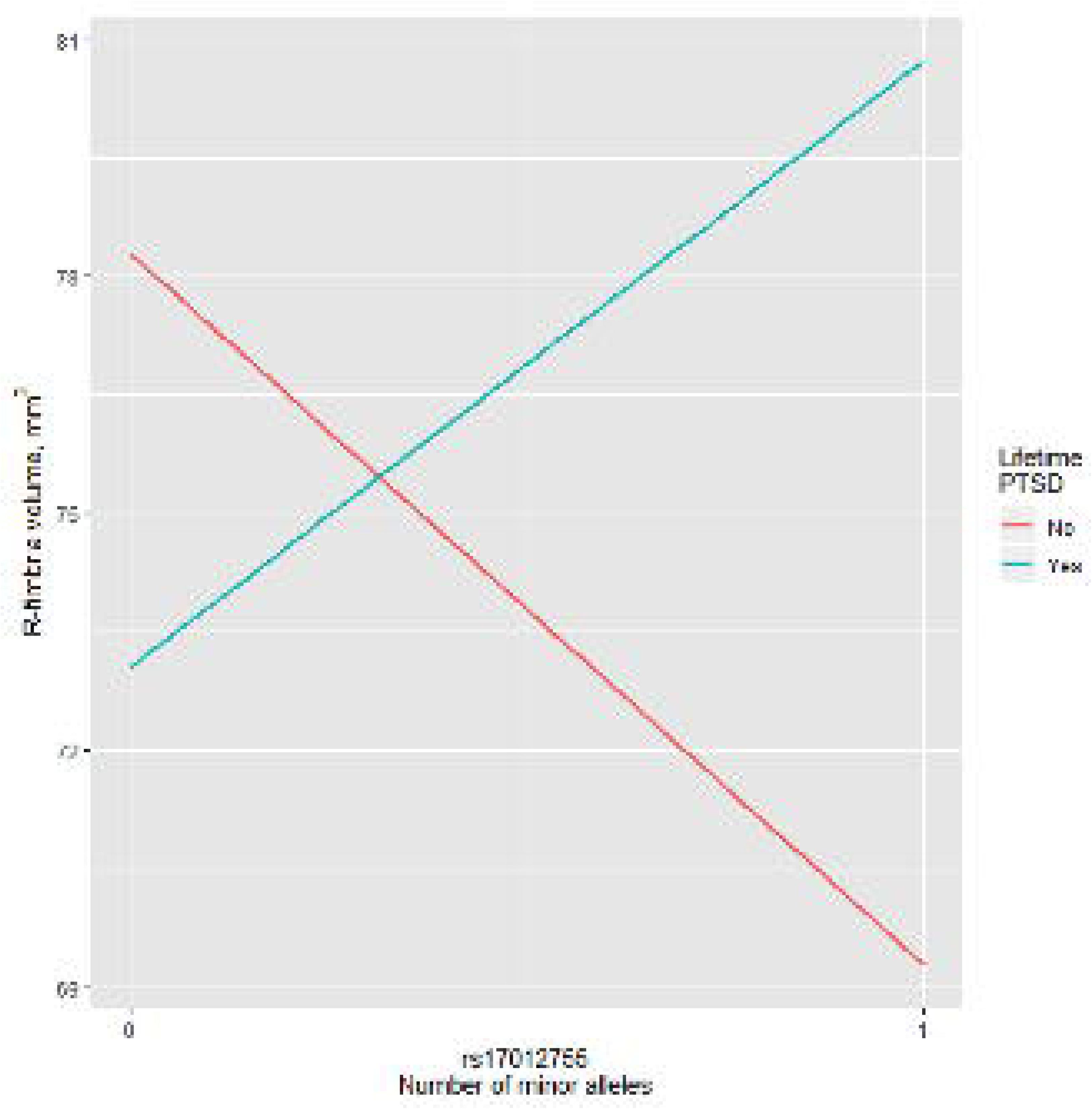

## DISCUSSION

We performed high-density GWAS of hippocampal subfields segmented by FreeSurfer 6.0 in trauma-exposed individuals with and without PTSD in a discovery sample of US military veterans and a replication sample of highly traumatized civilians. Several genetic markers reached genome wide significance for specific hippocampal subfields. Two SNPs that were involved in three genome-wide significant associations lie within genes(rs73008928 in *ARMC6* and rs7196581 in *RBBP6*). The SNP rs73008928 is particularly interesting as it has been identified as a methylation quantitative trait locus(QTL) and as an expression QTL, according to the ARIES(25) and GTEx databases(www.gtexportal.org)(26), respectively. Specifically, rs73008928 was found to be a trans-QTL for methylation probe cg08284873 in *ADARB2* using a sample of middle-aged individuals(**Table S1**)(27). RNA-editing deaminase-2(*ADARB2*) belongs to a family of genes whose other members edit RNAs encoding GluRs in the rat hippocampus(28). While not directly involved in RNA editing activity, *ADARB2* has been shown to inhibit activity of the other members of this gene family, suggesting that it plays a regulatory role in RNA editing(29). In addition to being a methylation QTL, rs73008928 is an expression QTL for *SUGP2* in several tissues, as well as for *TMEM161A* and *SLC25A42* in single tissues(**Table S2**). *SUGP2* is a splicing factor involved in pre-mRNA processing mechanisms(30); *TMEM161A*, when overexpressed, plays a role in protection against oxidative stress(31); and *SLC25A42* belongs to a large family of nuclear-encoded transporters that are involved in metabolic pathways and cell functions and is widely expressed in the central nervous system(32, 33). In summary, we have identified a SNP associated with R-HATA volume that also appears to impact RNA editing of GluRs and regulate expression of a gene involved in oxidative stress; both are processes which have been previously implicated in the pathophysiology of PTSD(34-36).

The other genome-wide significant SNP associated with hippocampal subfield volume is rs7196581 located in the *RBBP6* gene. This SNP was significantly associated with both L-CA1 volume and R-HATA volume. It falls within the promoter region of *RBBP6* and coincides with H3K27ac modification sites in several cell types, inclusive of neuronal progenitors, suggesting that this SNP functions as an enhancer(37). The CA1 has direct reciprocal projections to the medial prefrontal cortex(mPFC) that allow these regions to form a functional loop. This circuit enables interactions between cortical and subcortical areas during memory encoding and retrieval of episodic-like memories(38). Neurons in hippocampal CA1 code both space and time, allowing conjoint spatial and temporal representations of experiences(39). The critical role of these subfields in episodic memories makes them of particular interest in PTSD. Despite substantial research on purported episodic memory deficits in PTSD, which would implicate the functional loop, the evidence has been inconclusive(40). Severity of PTSD is negatively associated with mPFC activation elicited by material that participants have subsequently forgotten(41). Veterans with PTSD have lower activation in the hippocampus and amygdala when successfully encoding trauma-related stimuli into memory(42). Therefore, the present results that show an association of genetic markers with CA1 volume, but no interaction with PTSD diagnosis, may help build the current body of relevant knowledge.

All of the other SNPs associated with hippocampal subfields in this study were intergenic, indicating there may be some underlying regulatory genomic mechanism driving the statistical associations that we observed. Of interest, three significant SNPs reside in lincRNAs (rs6906714 in *LINC02571*, rs12880795 in *TUNAR*, and rs3811492 in *LINC01736*), which comprise the largest proportion of long non-coding RNAs(lnRNAs) in the human genome(43). While lncRNAs do not encode proteins, they and other non-coding RNAs, such as micro-RNAs(miRNAs), have been implicated in several neuropsychiatric disorders including PTSD(44-47). lncRNAs have also been associated with changes in gene expression in animal models of PTSD. For example, lncRNAs were differentially expressed in the hippocampus of rats exposed to stress-enhanced fear learning compared to control rats(48). In addition, RNA sequencing of mPFC in adult mice revealed changes in lncRNA expression after fear conditioning(49). The association we observed between rs12880795 in *TUNAR* and L-HATA volume was significantly replicated in the Grady Trauma Project cohort, providing additional confidence in this finding. These data suggest that the role of ncRNAs in PTSD should be studied further and be useful biomarkers or therapeutic targets.

The association between rs6906714 and R-fimbria volume was the most significant finding in this study. Notably, rs6906714 is also an expression QTL for several genes including *PSORS1C1* and *PSORS1C2*, which are psoriasis susceptibility genes that are expressed in thyroid tissue(**Table S2**). Genes associated with autoimmune disorders have been previously associated with PTSD(50); the overlap between stress disorders and immune/inflammatory disorders is an ongoing area of research(51). We also observed an interaction of rs6906714 with childhood trauma in the discovery cohort, and of rs6906714 with lifetime PTSD in the replication cohort, which is consistent with neuropsychiatric responses to stress that show increased risk of subsequent autoimmune disease(**Figure 2**)(51). The second strongest signal was between rs17012755, an intergenic SNP on chromosome 2, and R-fimbria volume. We also observed a significant interaction between this SNP and lifetime PTSD on R-fimbria volume in the replication cohort. Interestingly, in our larger PTSD cohort of over 2000 subjects, we observed nominal evidence for association between rs17012755 and PTSD(19).

The fimbria is a primary source of input to the dorsal hippocampus, which is a critical element in contextual fear conditioning. Therefore, contextual fear conditioning has become a widely adopted experimental model for studying PTSD(52, 53). Processing of contextual memories is mediated by robust projections from hippocampal neurons to the mPFC(54). Connectivity between the hippocampus and the mPFC, which involves the fimbria, are associated with mnemonic and emotion regulation deficits, which are consistent with the clinical presentation of PTSD(52). Recent evidence suggests that the mPFC directs the retrieval of context-appropriate episodic memories in the hippocampus(38, 55, 56). Electrolytic lesions to the fimbria in rats produce deficits in fear conditioning to contextual stimuli. The deficit in freezing, following dorsal hippocampus and fimbria lesions, is evident on both the conditioning day and the delayed extinction test(57). Indeed, contextual fear deficits in rats with hippocampal damage are equivalent following lesions to either fimbria, dorsal hippocampus, or entorhinal cortex(57). In addition to contextual fear conditioning, the fimbria plays an important role in spatial memory, which enables rats to learn the location of both a visible and hidden/submerged platform in the water maze task(58). However, rats with lesions to the fimbria learned to swim to the visible platform but were unable to navigate to the submerged platform in the same location(59,60).

The most significant effects in both discovery and replication cohorts involved R-fimbria volume. Deficits in contextual processing are considered to be representative of PTSD because intrusive memories and perceptions are experienced outside of the trauma context. Our finding that exposure to child trauma is a potent environmental exposure that interacts with genetic markers to influence brain structure is consistent with our recently published multi-cohort study in 1,868 subjects, which demonstrated childhood trauma exposure has a negative association with hippocampal volume but was not significant if PTSD was added as a covariate(3). Our finding underscores the uniquely deleterious role of trauma during childhood, which is a critical neurodevelopmental time period(61).

### Strengths and Limitations

The current work supports the value of imaging genetic studies. Our study was focused on hippocampal subfields, given its potential relevance to PTSD. We observed several strong associations, at least one of which demonstrated replication in an independent data set. However, our study is not without limitations. Due to the modest sample size of both the discovery and replication cohorts utilized, we lack statistical power to detect associations with small effect sizes of main effects as well as SNP x PTSD and SNP x childhood trauma interactions. As such, we restricted our investigation of interactions to only those SNPs which demonstrated a main effect with hippocampal subfield volume. In the near future, we plan to interrogate dramatically larger data sets accessed through worldwide consortia to ascertain these important interactions on a genome-wide scale(62).

One of the significant associations in our discovery cohort was confirmed in our replication cohort. However, the other nine discovery associations were not significant in the replication cohort. There are many reasons for this, the most likely being differences in population structure. The discovery cohort had nearly equal proportions of European Americans and African Americans, while the replication cohort was entirely African American. As such, the allele frequencies of the significant SNPs differed between the two cohorts, and in one case(rs2553974), the minor allele was reversed entirely. Other reasons for lack of replication could be cohort differences in trauma type and male/female ratio. The discovery cohort was comprised of military veterans while the replication cohort was entirely civilian, which could impact the ability to detect PTSD interactions. Nevertheless, we are encouraged that one association replicated despite differences between cohorts, which underscores the robustness of the underlying biological processes at play.

### Conclusion

Variation in the volume of fimbria, CA1, hippocampal amygdala transition area, and subiculum subfields is at least partially controlled by genetic factors as well as the interaction of genetics and environmental exposure to trauma. These findings provide new avenues for understanding neural circuits that are implicated in contextual fear conditioning and other established behavioral models of PTSD. Ultimately the promise of finding genetic determinants of PTSD is that they signal the presence of etiologic pathways for targeted. Attempts to find disease-associated genetic variation that point to molecular mechanisms of pathogenesis has proven challenging due to the polygenicity of clinical phenotypes(63, 64). Leveraging neuroimaging phenotypes may offer a shortcut over clinical phenotypes in identifying these elusive genetic markers and relevant neurobiological pathways(62, 65).

## ACKNOWLEDGMENTS

The views expressed in this manuscript are those of the authors and do not necessarily represent the views of the funding agencies or the federal government. This research was supported by the U.S. Department of Veterans Affairs (VA) Mid-Atlantic Mental Illness Research, Education, and Clinical Center (MIRECC) core funds. Dr. Morey also received financial support from the US Department of Veterans Affairs (VA) Office of Research and Development (5I01CX000748-01, 5I01CX000120-02). Additional financial support was provided by the National Institute for Neurological Disorders and Stroke (R01NS086885-01A1). Dr. Dedert is funded by a VA Clinical Science Research and Development (CSR&D) Career Development Award (CDA) (IK2CX000718). Dr. Naylor is funded by a VA Rehabilitation Research and Development (RR&D) CDA (1lK2RX000908). Dr. Van Voorhees is funded by a Department of Veterans Affairs Rehabilitation Research and Development Career Development Award (1K2RX001298).Dr. Kimbrel is funded by a Department of Veterans Affairs Clinical Science Research and Development Career Development Award (IK2CX000525). Dr. Beckham also received financial support from the U.S. Department of Veterans Affairs (VA) Office of Research and Development (1IK6CX001494). We thank the willing participation of Veterans in this research.

